# Synchrony and perturbation transmission in trophic metacommunities

**DOI:** 10.1101/2020.01.21.914093

**Authors:** Pierre Quévreux, Matthieu Barbier, Michel Loreau

## Abstract

In a world where natural habitats are ever more fragmented, the dynamics of meta-communities is essential to properly understand species responses to perturbations. If species’ populations fluctuate asynchronously, the risk of their simultaneous extinction is low, thus reducing the species’ regional extinction risk. We propose a metacommunity model consisting of two food chains connected by dispersal to study the transmission of small perturbations affecting populations in the vicinity of an equilibrium. We show that perturbing a species in one patch can lead to asynchrony between patches if the perturbed species is not the most affected by dispersal. Dispersal affects rare species the most, thus making biomass distribution critical to understand the response of trophic metacommunities to perturbations. We further partition the effect of each perturbation on species synchrony when several independent perturbations are applied. Our approach allows disentangling and predicting the responses of simple trophic metacommunities to perturbations, thus providing a theoretical foundation for future studies considering more complex spatial ecological systems.

## Introduction

Biodiversity is under increasing anthropic perturbations that alter populations and community dynamics (see the last IPBES assessment (Díaz et al., 2019)). In particular, communities live in more and more fragmented habitats (Haddad et al., 2015), which reduce dispersal and partially isolate populations. The metacommunity framework is key to address the responses of species and communities to perturbations in this changing world (Leibold et al., 2004; Amarasekare, 2008; Leibold and Chase, 2017). Small isolated populations are more prone to extinction (Purvis et al., 2000), and simultaneous local extinctions in all localities lead to a global extinction. Thus, the asynchrony between different populations of the same species is a fundamental mechanism ensuring the persistence and temporal stability of an entire metapopulation at the landscape scale as it reduces the species’ extinction risk in all patches simultaneously (Blasius et al., 1999).

While dispersal tends to synchronise populations of the same species (Abbott, 2011), dispersal of specific trophic levels can lead to synchrony or asynchrony between the dynamics of the various species in food chain models (Koelle and Vandermeer, 2005; Pedersen et al., 2016). Species that disperse or forage across several communities can propagate trophic cascades in space, as shown empirically and theoretically (Knight et al., 2005; McCoy et al., 2009; Casini et al., 2012; García-Callejas et al., 2019); depending on which trophic levels disperses the strength of trophic cascades within each community can be amplified or dampened (Leroux and Loreau, 2008). In addition, different food chain lengths in each locality can lead to opposite responses of different populations of the same species (Wollrab et al., 2012).

The dispersal of top predators has been particularly studied as generalist consumers linking different food webs by feeding on multiple energetic channels are ubiquitous across ecosystems (Rooney et al., 2006, 2008; Wolkovich et al., 2014; Ward et al., 2015). In particular, mobile predatory fish couple pelagic and benthic compartments in aquatic ecosystems (Vander Zanden and Vadeboncoeur, 2002; Vadeboncoeur et al., 2005), and predator dispersal leads to trophic cascade in surrounding ecosystems (Knight et al., 2005; Casini et al., 2012; Tscharntke et al., 2012).

Theoretical models suggest that asymmetry between the coupled food chains is key: mobile top predators can stabilise biomass dynamics by promoting asynchrony between the different populations of the consumed species if one of them is preferred (McCann et al., 1998) or if the energy transfer is faster in one community (Rooney et al., 2006). Asymmetry promotes asynchrony even when top predator populations are under correlated environmental perturbations (Vasseur and Fox, 2007). In contrast, predators tend to synchronise their prey populations in the absence of asymmetry between communities (Koelle and Vandermeer, 2005).

Many of these theoretical studies have considered the synchrony between food chains displaying chaotic dynamics or limit cycles (McCann et al., 1998; Koelle and Vandermeer, 2005; Rooney et al., 2006), which are characteristic of strong top-down control (Barbier and Loreau, 2019). Unfortunately, many of the mechanisms cited (*e.g.* top predator coupling or asymmetry) act simultaneously and interact with the variability generated by the limit cycles of food chain dynamics. Variability can also be generated by stochastic external perturbations, but few studies have considered these (Vasseur and Fox, 2007) while Wang et al. (2015) used them successfully to investigate the stability of competitive metacommunities.

Here we propose a first step toward a more systematic approach to synchrony in trophic metacommunities. The aim of our contribution is to understand what shapes synchrony in a broad spectrum of ecological settings, dominated by either bottom-up or top-down control (Barbier and Loreau, 2019) within a food chain, and by either trophic or spatial mechanisms at each trophic level. The relative importance of local dynamics and dispersal is what distinguishes different metacommunity paradigms (Leibold et al., 2004; Leibold and Chase, 2017). It also controls different recovery regimes after perturbations. For instance, Zelnik et al. (2019) showed that, with low dispersal and fast local dynamics, the system recovers locally from the perturbation, while with high dispersal and slow local dynamics, perturbations propagate across the whole system. A synthetic understanding of synchrony may thus be achieved by quantifying the propagation of perturbations, both vertically along food chains, and horizontally across space.

This approach allows us to identify key ecological factors that affect synchrony. In a single food chain, Barbier and Loreau (2019) showed that a few parameters control the biomass distribution among trophic levels (*i.e.* top or bottom-heavy pyramids, trophic cascades) and the overall top-down or bottom-up behaviour of the system. In turn, the biomass distribution drives many processes in food web dynamics. For instance, Arnoldi et al. (2019) showed that stochastic perturbations can affect species more or less depending on their relative abundance. In addition, biomass distribution determines the relative importance of local dynamics and dispersal, as we found that less abundant species are more affected by dispersal. As noted above, the higher dispersal of particular trophic levels has strong consequences for food web dynamics (Koelle and Vandermeer, 2005; Pedersen et al., 2016). Therefore, if a perturbation does not affect the species that disperse the most, its transmission through the food chain and between communities can lead to different responses in different localities, and hence generate asynchrony between them.

We develop a model of coupled food chains based on these recent studies (1) to understand the role of biomass distribution in the relative importance of local demographic processes and dispersal and (2) to disentangle the effects of multiple perturbations on the overall synchrony between populations at the same or different trophic levels. As a starting point, we consider a simple setting with Lotka-Volterra dynamics and stochastic external perturbations around an equilibrium. This allows us to partition the variability and the correlations generated by multiple perturbations. Partitioning approaches provide a powerful way to disentangle the effects of different mechanisms and to assess their relative importance (Price, 1970; Loreau and Hector, 2001; Jaillard et al., 2018). It also allows us to use simple scenarios in which a single species is perturbed as building blocks to understand more complex systems with multiple perturbations. Thus, we could assess the contribution of each species and their influence on other species to explain the synchrony or the asynchrony between the different populations.

## Material and methods

### The metacommunity model

We extend the model developed by (Barbier and Loreau, 2019). They considered a food chain model with four trophic levels and a simple metabolic parametrisation, for which they described biomass repartitions (Fig.1A) and responses to perturbations. Their model corresponds to the “intra-patch dynamics” part of equation (1) to which we graft a dispersal term to consider a metacommunity with two patches

**Figure 1:**
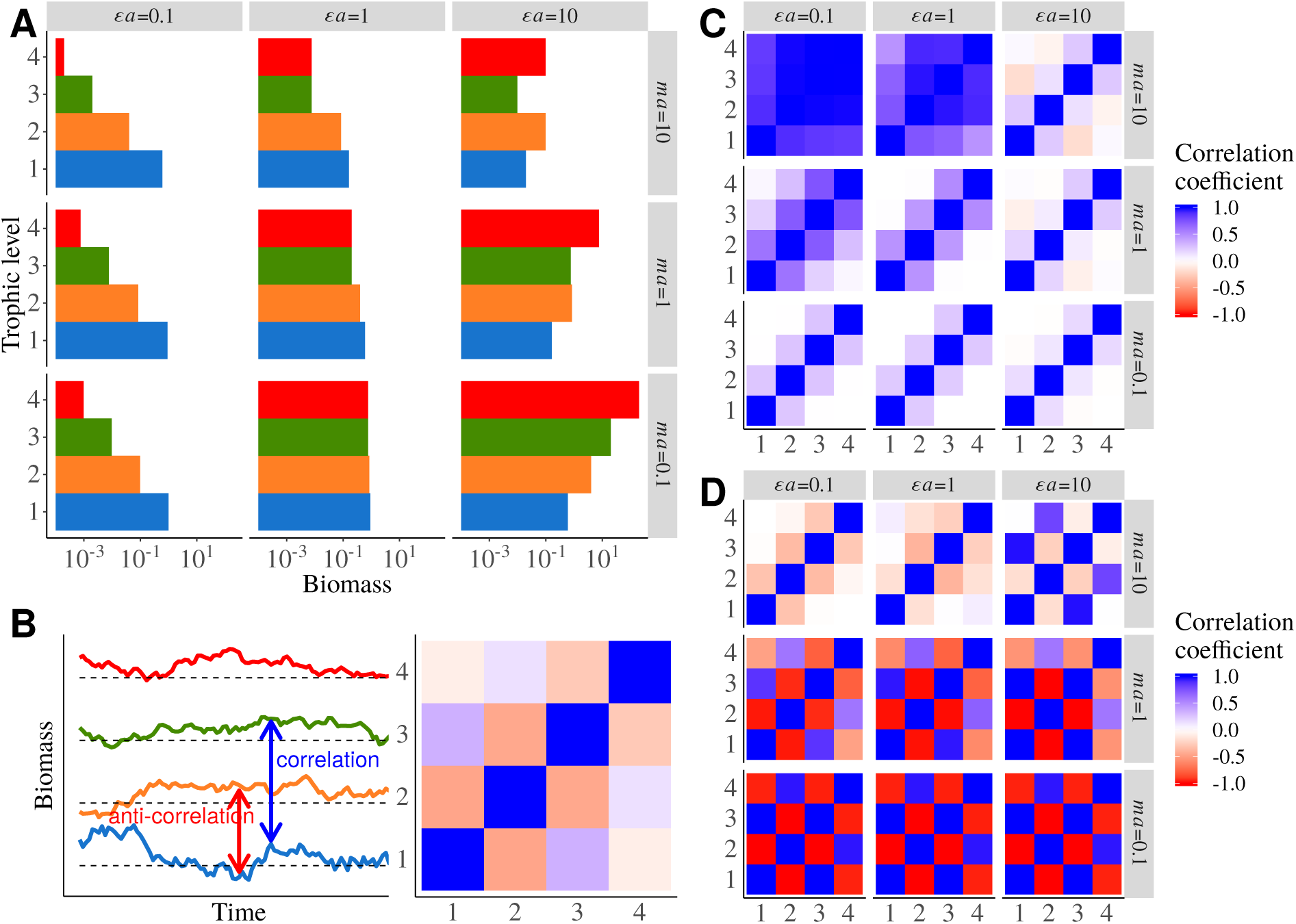
General description of an isolated food chain (*d*_*i*_ = 0, no dispersal) for nine combination of the physiological and ecological parameters *ϵa* and *ma* that respectively describe the positive effect of biomass consumption and the negative effects mortality due to predation (see Barbier and Loreau (2019)). **A)** Biomass distribution among trophic levels. **B)** Correlation between species biomass dynamics. The correlations seen in time series are represented by a correlation matrix where each element is the correlation coefficient between two species. Thus, the matrix is symmetric and the diagonal elements are equal to 1 as each species is perfectly autocorrelated. **C)** Correlation matrix within a food chain with a demographic stochastic perturbation applied to primary producers. **D)** Same correlation matrix with a demographic stochastic perturbation applied to top predators.

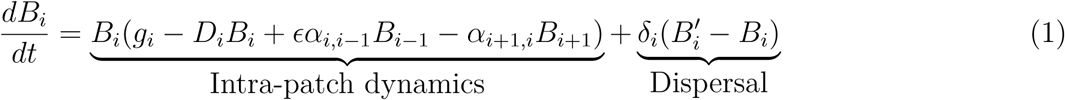

where *B*_*i*_ is the biomass of trophic level *i* in the patch of interest and 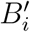 its biomass in the other patch, *ϵ* the biomass conversion efficiency and *α*_*i,j*_ the interaction strength between consumer *i* and prey *j*. Species *i* disperses between the two patches at rate *δ*_*i*_. The density independent growth or mortality rate *g*_*i*_ and the density dependent mortality rate *D*_*i*_ scale with species metabolic rates *m*_*i*_:

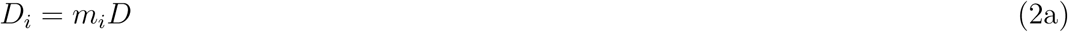

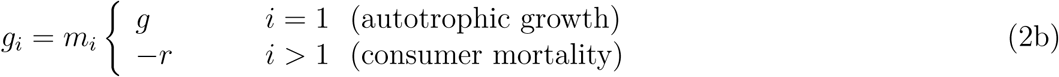

The predator-prey metabolic rate ratio and the interaction strength to self-regulation ratio are assumed to be constant:

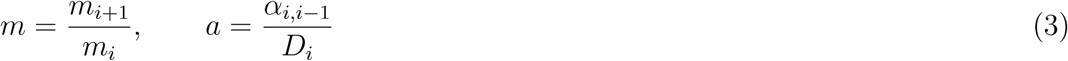

We also defines *d*_*i*_ the dispersal rate relative to self-regulation:

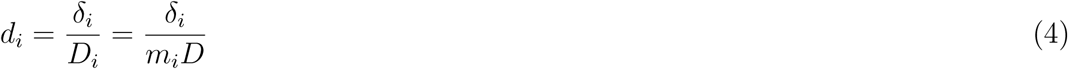

and the time scale of the system is defined by setting the metabolic rate of the primary producer to unity. Thus, we can transform the equations for primary producers (*i* = 1):

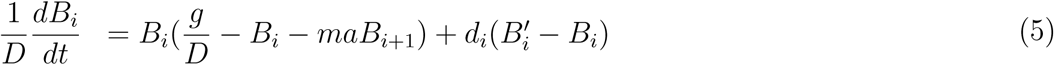

and consumers:

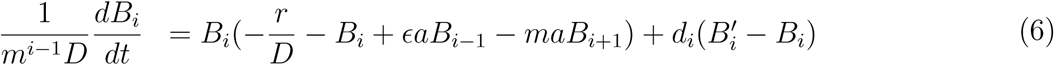

See Table 1 summarising parameter definition and values.

**Table 1:** Table of parameters. Only combinations of *m* and *a* leading to the desired values of *ma* were kept.

| parameter | interpretation | value |
| --- | --- | --- |
| $\sigma$ | intensity of stochastic noise | $\{10^{-3}, 10^{-5}\}$ |
| $g$ | net growth rate of primary producers | 1 |
| $r$ | death rate of consumers | 0 |
| $D$ | self regulation | 1 |
| $\epsilon$ | conversion efficiency | 0.65 |
| $m$ | predator/prey metabolic rate ratio | $\{0.0065, 0.065, 0.65, 6.5, 65\}$ |
| $a$ | attack rate | $\{1/6.5, 1/0.65, 1/0.065\}$ |
| $\epsilon a$ | positive effect of prey on predators | $\{0.1, 1, 10\}$ |
| $ma$ | negative effect of predators on prey | $\{0.1, 1, 10\}$ |
| $d_i$ | dispersal rate | $[10^{-5}, 10^5]$ |

### Biomass at equilibrium

The system of equations (5) and (6) at equilibrium cannot be solved analytically. We used the multidimensional root-finder algorithm of the GNU Scientific Library version 2.5 (Galassi, 2009) initialised with the equilibrium biomass of the system without dispersal that can be easily calculated analytically, by solving the following equations:

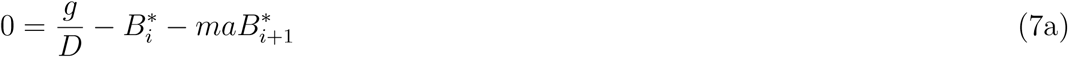

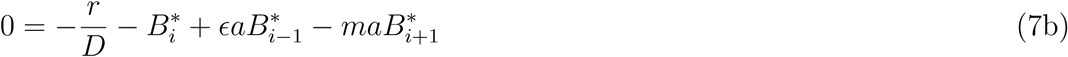

This can be expressed as a matrix product:

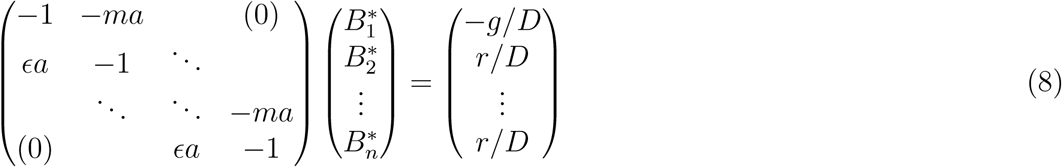

solved by the tridiagonal solver algorithm of the GNU Scientific Library version 2.5 (Galassi, 2009).

### Response to perturbations

#### Stochastic perturbation in the vicinity of equilibrium

The general system with *S* species is defined by:

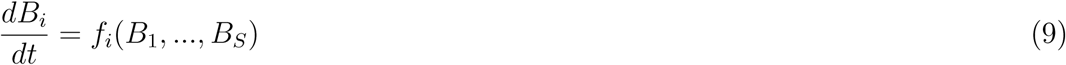

*B*\* defines the equilibrium at which the community matrix (or Jacobian matrix) *J* is defined by:

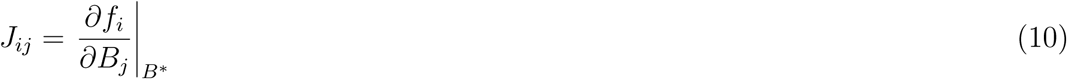

The system can be linearised in the vicinity of *B*\*:

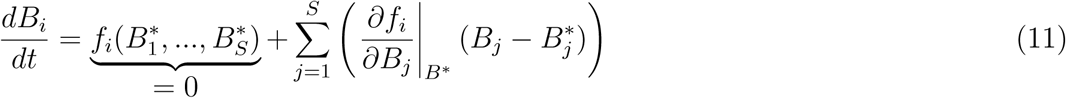

Thus, by setting 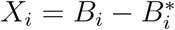 the deviation from equilibrium, we have:

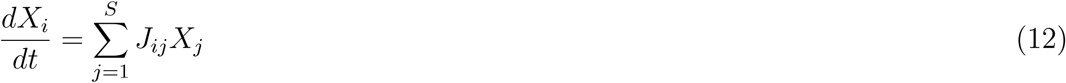

Then, we can consider a small perturbation due to *R* factors defined by 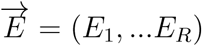 whose effects on 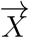 are defined by the matrix *T* (Arnoldi et al., 2016). We have the equation:

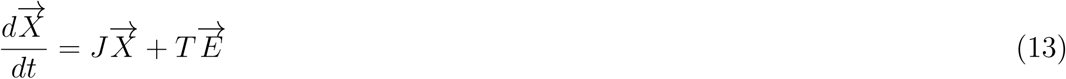

The elements of 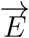 are defined by stochastic perturbations *E*_*i*_ = *σ*_*i*_*dW*_*i*_ with *σ*_*i*_ their standard deviation and *dW*_*i*_ a white noise term with mean 0 and variance 1. In addition, perturbations scale with each species biomass 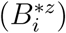. We can consider three types of perturbation (Haegeman and Loreau, 2011; Arnoldi et al., 2019): exogenous if *z* = 0 (no scaling with biomass), demographic if *z* = 0.5 and environmental if *z* = 1. We chose the demographic perturbation as it has no bias for rare or abundant species (Arnoldi et al., 2019). In our case, *T* contains three block diagonal matrix corresponding to each type of perturbation as species can receive several types of perturbations (see Fig.A-5 in the supporting information):

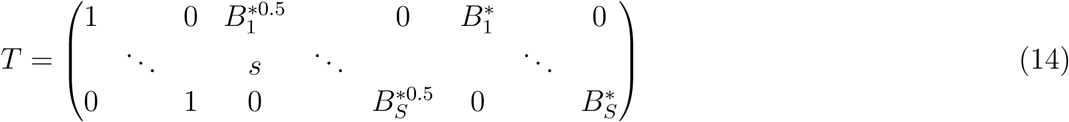

#### Computing the covariance matrix

The stationary covariance of 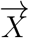 at equilibrium is defined by *C*\* = 𝔼(*X*\**X*\*^⊤^) and is the solution of the Lyapunov equation (Arnold, 1974; Wang et al., 2015; Arnoldi et al., 2016; Shanafelt and Loreau, 2018):

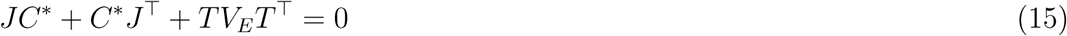

with *J* the Jacobian matrix, *V*_*E*_ the covariance matrix of stochastic perturbations whose diagonal elements equal to 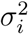 (and non-diagonal elements equal to zero if perturbations are independent) and *T* the matrix defining how the perturbations apply to the system (see equation (13)). *C*\* can be calculated using a *Kronecker product* (Nip et al., 2013). The *Kronecker product* of an *m* × *n* matrix *A* and a *p* × *q* matrix *B* denoted *A* ⊗ *B* is the *mp* × *nq* block matrix given by:

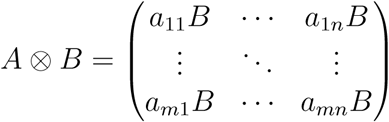

We define 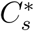 and (*TV*_*E*_*T*^⊤^)_*s*_ the vectors stacking the columns of *C*\* and *TV*_*E*_*T*^⊤^ respectively. Thus, equation (15) can be rewrite as:

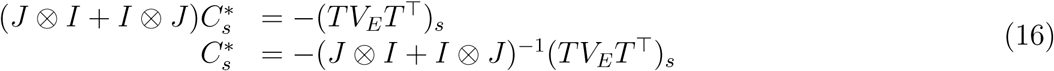

From the covariance matrix *C*\* whose elements are *w*_*ij*_ we can compute the correlation matrix *R*\* of the system whose elements *ρ*_*ij*_ are defined by:

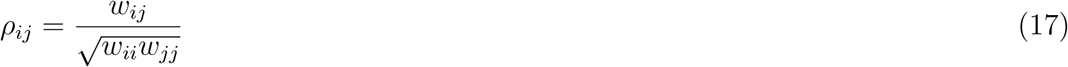

#### Local demography versus dispersal processes

Correlation between different populations of the same species depends on the relative importance of demographic and dispersal processes: dispersal processes tend to correlate (or anti-correlate) populations while demographic process tend to decorrelate them. We define a metric *M*_1_ that describes the relative weight of these two processes by taking the absolute values of the each element of equation (1) to assess the sheer intensity of local demographic processes and dispersal processes:

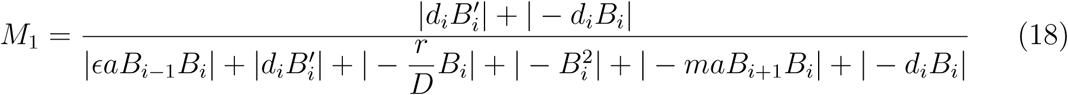

### Parameters

As in Barbier and Loreau (2019), we explore the response of our model over two order of magnitude of the aggregated parameters *ϵa* and *ma* as they allow to explore all the distinct qualitative behaviours of an isolated food chain (see Fig1A for the biomass repartition for instance). See table B-2 in the supporting information displaying the values of *m* and *a* used to get each desired value of *ϵa* and *ma*.

## Results

### General responses of the food chain model to perturbations

We first describe the biomass distribution and the responses to perturbations of an isolated food chain (*i.e.* without considering spacial dynamics). We use a broad range of physiological and ecological parameters to describe all the possible responses of the food chain model (Fig.1A). *ma* represents the strength of negative interactions (mortality due to predation) while *ϵa* represents the strength of positive interactions (biomass gain due to consumption). As in Barbier and Loreau (2019), the food chain displays various biomass distributions in different regimes: bottom-heavy (for *ϵa* = 0.1) biomass pyramids, top-heavy biomass pyramids (for *ϵa* = 10 and *ma* = 0.1) or alternating “cascade” patterns (for *ϵa* = 10 and *ma* = 10).

In each case, we can capture the dynamical behaviour of the food chain by considering the correlation matrix of the response of each species to perturbations applied to specific trophic levels (Fig.1B). Perturbing primary producers leads to bottom-up responses in which adjacent trophic levels are correlated, *i.e.* their biomasses respond in the same way, (Fig.1B and 1C) while perturbing top predators leads to top-down responses in which adjacent trophic levels are anti-correlated, *i.e.* their biomasses respond in opposite ways, (Fig.1B and 1D). When all species receive independent stochastic demographic perturbations (Fig.A-1A in the supporting information), the correlation pattern is dominated by bottom-up effects for high values of *ma* (*ma* = 10, which corresponds to the strongest responses in Fig.1C) and is top-down for low values of *ma* (*ma* ≤ 1, which corresponds to the strongest responses in Fig.1D, see also Fig.A-1C in the supporting information).

### Propagation of a perturbation when one species disperses

Perturbations can propagate vertically within a food chain or horizontally between food chains. To understand how these two types of propagations shape the synchrony between patches we first consider a simple case where only primary producers are perturbed in patch #1 (patch #2 being the unperturbed patch) and only top predators disperse (Fig.2A). In patch #1, the perturbation has a bottom-up transmission that correlates species (Fig.2B, label (1)) as in Fig.1C where primary producers are also directly perturbed. While in patch #2, the perturbation has a top-down transmission (Fig.2A), leading to an anti-correlation of adjacent trophic levels (Fig.2B, label (2)), which is similar to Fig.1D as the transmission of the perturbation by top predators is equivalent to a direct perturbations of top predators in patch #2. Then, the different correlation patterns within each patch affect the synchrony between the two patches. First, the two populations of top predators are perfectly correlated as they are directly coupled through dispersal (Fig.2B, label (3)). Second, the populations of carnivores are anti-correlated because they are respectively correlated and anti-correlated to top predators in patch #1 and #2 (Fig.2B, (4)). Similarly, the correlation between each trophic level and top predators in each patch drives the correlation between the two population at lower trophic levels.

**Figure 2:**
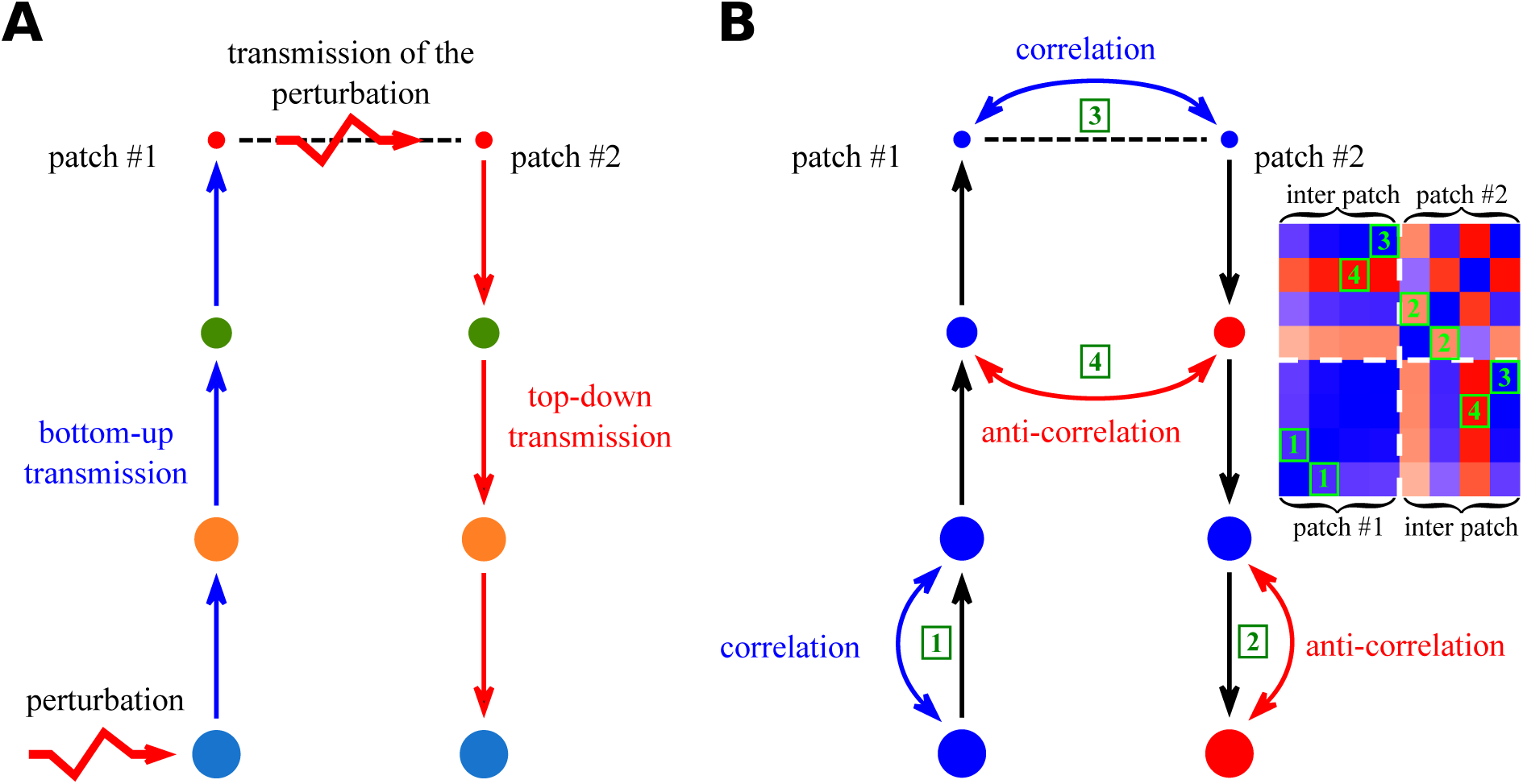
Transmission of perturbations between the two patches (primary producers perturbed in patch #1, *ϵa* = 0.1, *ma* = 10 and only top predators disperse). Disk size represents species abundance. **A)** Only top predators are able to disperse, transmitting the perturbation between patches. They convert the bottom-up perturbation from patch #1 into a top-down perturbation in patch #2. **B)** The bottom-up transmission in patch #1 leads to correlations between adjacent trophic levels (label 1) while the top-down transmission leads to anti-correlations between adjacent trophic levels in patch #2 (2). Dispersal directly couples the two populations of top predators that act as one unique population, thus, they are completely correlated (3). The different correlation patterns within each patch lead to correlations or anti-correlations between populations of the same species in different patches depending on its distance from top predators (4).

### For whom does dispersal matter?

Now, all species disperse at the same rate *d*_*i*_ but we still consider perturbations only affecting the primary producers in patch #1. Even if all dispersal rates are equal, the relative importance of dispersal processes compared to intra-patch demography quantified by *M*_1_ (see equation (18)) differs between species. When dispersal rates *d*_*i*_ increase, *M*_1_ first increases for top predators, then for carnivores and so on until primary producers (Fig.3E). This is due to biomass distribution (Fig.1A) as dispersal scales linearly with biomass while intra-patch demography scales with squared biomass (self-regulation) or biomass products (predation) (see equation (6) and Fig.A-1E in the supporting information).

**Figure 3:**
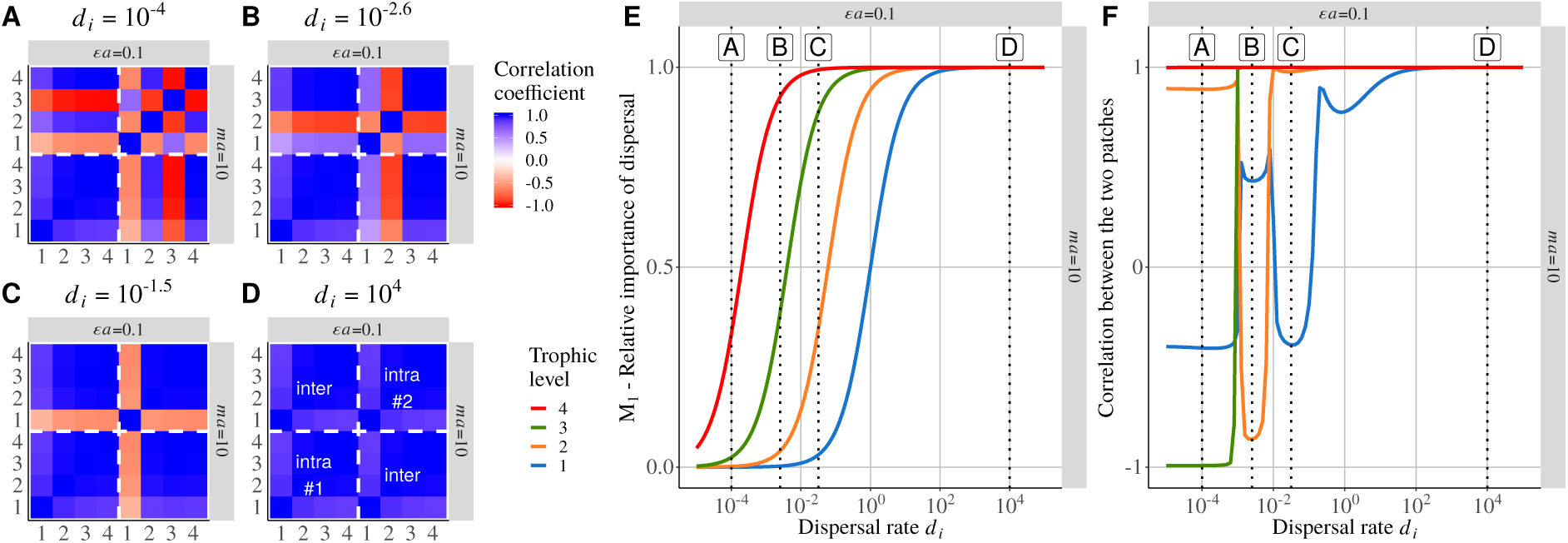
Correlation between populations of four species forming a food chain present in two connected patches (*ϵa* = 0.1, *ma* = 10). Primary producers in patch #1 receive demo-graphic perturbations while patch #2 is not directly perturbed. **A), B), C)** and **D)** are correlation matrix between species within and between patches for four different dispersal rates. Diagonal blocks represent intra-patch species correlations while the other blocks represent inter-patch species correlations (see the labels in **D)**). The bottom-left block represents the perturbed patch (#1) while the top-right block represents the unperturbed patch (#2) (see Fig.2B). **E)** *M*_1_, ratio of dispersal processes to the sum demographic and dispersal processes (see equation (18)) for each trophic level with increasing dispersal rates. Labels A, B, C and D respectively refer to the values of dispersal rates used to plot the correlation matrix presented in **A), B), C)** and **D). F)** Correlation between populations of the same species from two patches for increasing dispersal rates *d*_*i*_ (equal for all species). The represented correlations are equal to the diagonal elements of the off-diagonal blocks of correlation matrix.

At low dispersal rates (*e.g. d*_*i*_ = 10^−4^), dispersal matters only for top predators (Fig.3E label A), leading to a situation already described by Fig.2. At intermediate dispersal rates (*e.g. d*_*i*_ = 10^−2.6^), dispersal also matters for carnivores (Fig.3E label B). Thus, top predators and carnivores are correlated between patches and we observe anti-correlations between adjacent trophic levels lower than 4 (Fig.3B). This time, this leads to the anti-correlation of sub-populations of herbivores (Fig.3F label B) while they were correlated previously (Fig.2B and Fig.3F label A). Therefore, each time dispersal starts to matter for another trophic level, the correlation pattern in patch #2 changes (Fig.3A-D), leading to shifts between correlations and anti-correlations between the populations of lower trophic levels (Fig.3F).

### Multiple perturbation partitioning

The case displayed in Fig.3 was easy to handle as only one perturbation was applied and we knew for which species dispersal mattered. Such a simple case can actually act as a building block to understand correlation patterns produced by multiple perturbations. In fact, for *R* independent perturbations, the covariance matrix 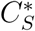 is equal to the sum of the covariance matrices 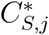 obtained when only one perturbation *j* is applied, 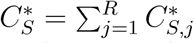 (see section Appendix A in the supporting information). Then, correlations between the populations of species *i* can be expressed as the sum of the correlations obtained when each perturbation *j* is applied alone weighted by the corresponding variance in the two patches.

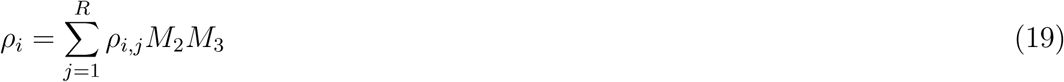

*R* is the number of independent perturbations, *ρ*_*i*_ is the correlation coefficient between the two populations of species *i* and *ρ*_*i,j*_ is the same correlation coefficient in the case where only perturbation *j* is applied. *M*_2_ quantifies the variability generated locally by perturbation *j* that is effectively transmitted to the other patch (Fig.4A). If *M*_2_ is close to zero, the perturbation is poorly transmitted and the two patches will probably be asynchronous. *M*_3_ weights each *ρ*_*i,j*_ by the variability generated by perturbation *j* compared to the other perturbations (Fig.4B). If *M*_3_ is low, perturbation *j* would generate less variability than the other perturbations and the associated correlation *ρ*_*i,j*_ will not significantly contribute to the correlation *ρ*_*i*_ generated by all perturbations.

**Figure 4:**
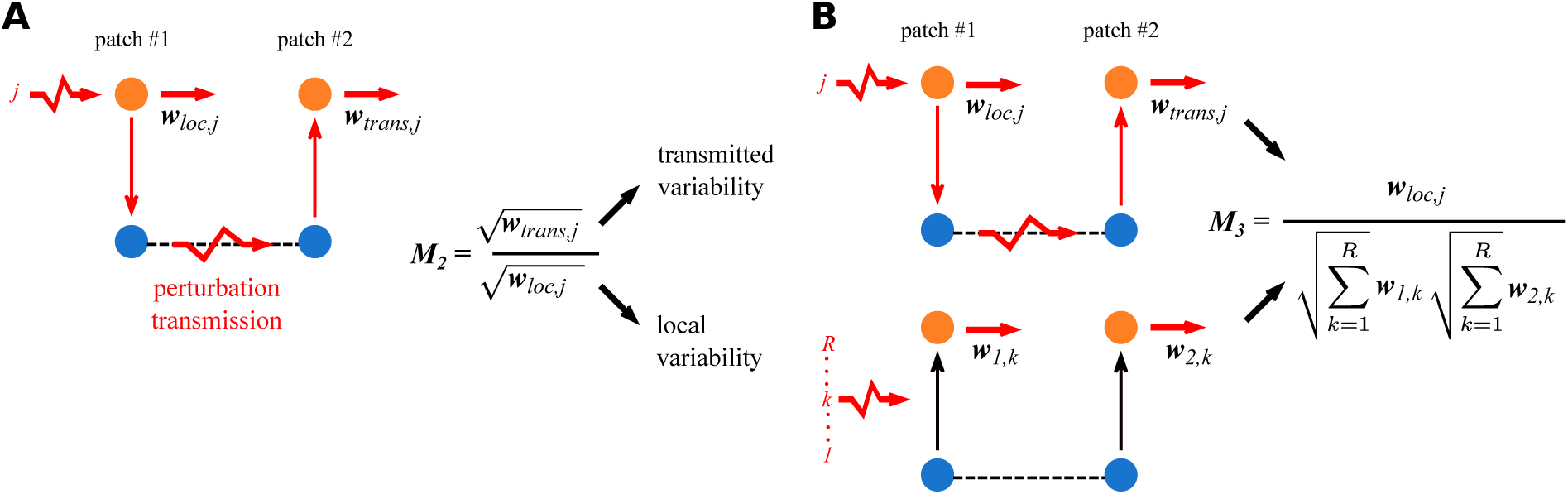
Metrics weighting the contribution of each perturbation to the correlation pattern generated by multiple perturbations. *w*_1,*j*_ and *w*_2,*j*_, which are element of the matrix 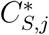, are the variance of species *i* in patches #1 and #2 respectively when perturbation *j* is applied. We define *w*_*loc,j*_ the variability directly generated by the perturbation and *w*_*trans,j*_ the variability transmitted in the other patch. *w*_*loc,j*_ and *w*_*trans,j*_ are respectively equal to *w*_1,*j*_ (or *w*_2,*j*_) and *w*_2,*j*_ (or *w*_1,*j*_) when perturbation *j* is applied in patch #1 (or #2). **A)** *M*_2_ is the ratio of transmitted variability *w*_*trans,j*_ to local variability *w*_*loc,j*_. **B)** *M*_3_ weights the effect of each perturbation *j* by the variability it generates locally compared to the other perturbations.

In the following, we present a simple case with two species in each patch receiving independent demographic stochastic perturbations and only primary producers able to disperse (see Fig.A-3 in the supporting information for an example with four species).

In Fig.5, we illustrate step by step the decomposition of the correlation pattern generated by multiple perturbations (Fig.5G). When only primary producers are perturbed in patch #1, both primary producers and herbivores are correlated due to the bottom-up transmission of the perturbation in both patches as only primary producers disperse (Fig.5A). However, when only herbivores are perturbed, herbivores become decorrelated as dispersal rates *d*_*i*_ increase (Fig.5B) due to the weak correlation between adjacent trophic levels for *ϵa* = 10 and *ma* = 10 (see Fig.1C and 1D).

**Figure 5:**
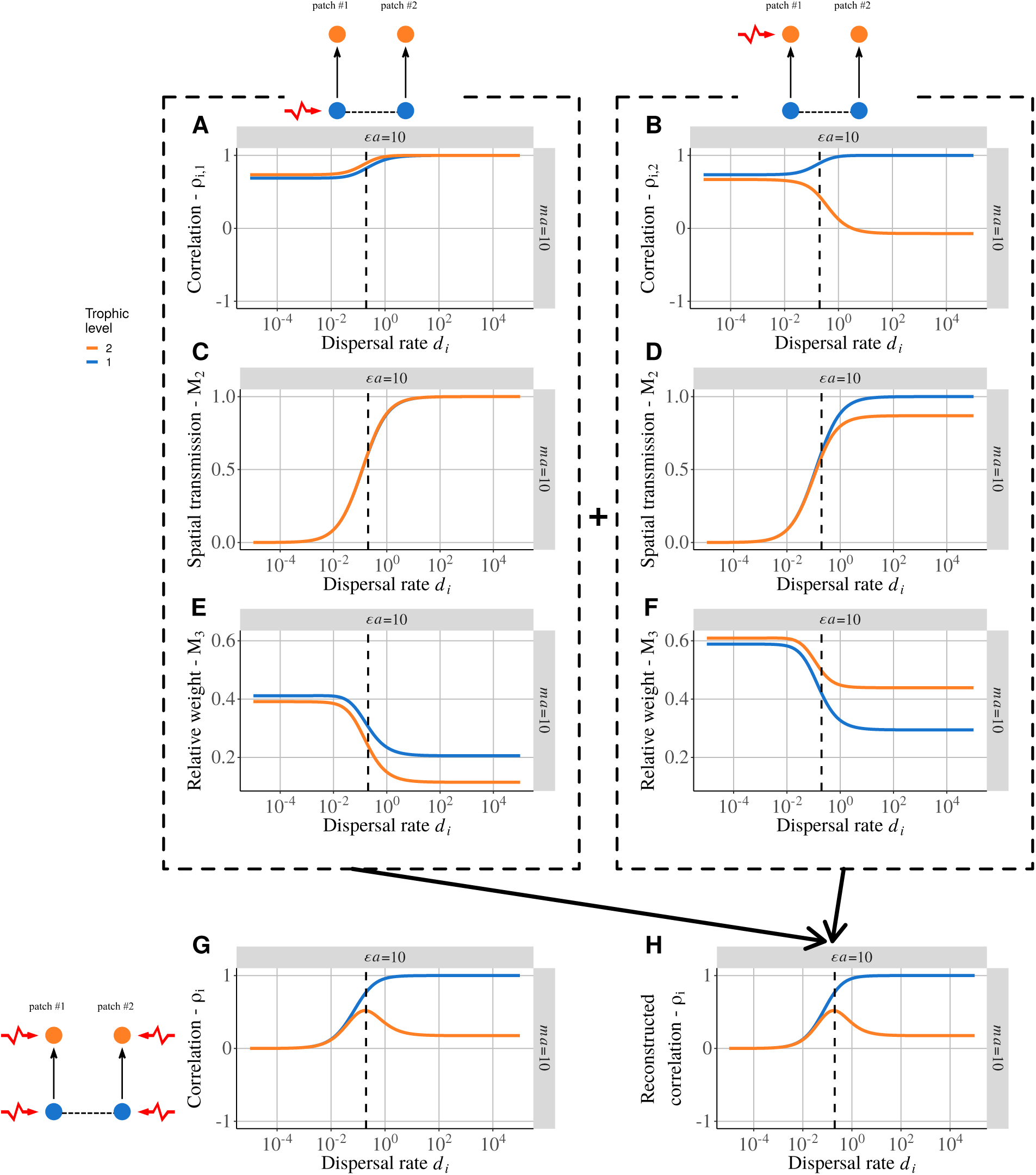
Detailed correlation pattern between two coupled plant-herbivore food chains for *ϵa* = 10 and *ma* = 10 with increasing dispersal rates *d*_*i*_. Only primary producers are able to disperse. **A)** Correlation between patches when only primary producers and **B)** herbivores from patch #1 are perturbed. **C)** Relative importance of local transmitted variability to local variability (*M*_2_) when primary producers and **D)** when herbivores are perturbed in patch #1. **E)** Relative weight of the variance generated by each perturbation (*M*_3_) when primary producers and **F)** when herbivores are perturbed in patch #1. **G)** Correlation between patches when independent demographic stochastic perturbations are applied to all species of each patch. **H)** Reconstructed correlation pattern obtained thanks to equation (19). **H**=2(**A**×**C**×**E**+**B**×**D**×**F**) by symmetry as both patch #1 and #2 receive similar independent perturbations.

Our metric *M*_2_ is equal to zero at low dispersal rates *d*_*i*_ (Fig.5C and 5D), thus indicating that the perturbations in patch #1 are weakly transmitted in patch #2. At high dispersal rates *d*_*i*_, *M*_2_ tends to 1 as species become perfectly correlated except for herbivores in Fig.5D. In this case, as they are perturbed but do not disperse, the perturbation is attenuated during its transmission through primary producers.

In this example, our metric *M*_3_ is higher for both primary producers and herbivores when the perturbation is applied to herbivores (Fig.5F) than to primary producers (Fig.5E). This means that perturbations applied to herbivores generate most of the variability in the metacommunity and the correlation pattern in Fig.5B thus strongly contributes to the reconstructed correlation pattern gathering the effects of all perturbations (Fig.5H) following equation (19).

Now we have the response of all the elements of equation (19), we can explain the correlation pattern seen in Fig.5H. At low dispersal rates *d*_*i*_, perturbations are not transmitted (*M*_2_ = 0), letting the two patch independent and uncorrelated, while at high dispersal rates *d*_*i*_, the correlation pattern is similar to Fig.5B as herbivore perturbation generates most of the variability. In between, we have a humped-shaped relationship between herbivore population correlation and dispersal rates *d*_*i*_ because when perturbations start to be transmitted (Fig.5C and 5D), herbivore populations are correlated (left to the dashed line) (Fig.5A and 5B). Then, the decrease in Fig.5B leads to the decrease seen in Fig.5H. The reconstructed correlation pattern in Fig.5H is identical to the correlation pattern obtained by perturbing directly each species in each patch (Fig.5G), thus demonstrating the validity of equation (19) (see Fig.A-4 in the supporting information).

## Discussion

Our metacommunity model aimed to understand how perturbations propagates vertically in patches and horizontally between patches to identify under which conditions species responses in different patches can be synchronous or asynchronous. First, we found that less abundant species are more affected by dispersal. Thus, even when all species disperse at the same rate, the biomass distribution in a food chain determines for which species dispersal contributes most to biomass dynamics. In addition, if the perturbed species does not disperse enough to synchronise its different populations, the perturbation can be transmitted by other species. In such a situation, we found that species responses in different patches can be asynchronous. Second, we found that the effects of multiple independent perturbations can be partitioned. This enabled us to use simple situations in which a single species is perturbed as building blocks to analyse more complex systems with multiple perturbations. Thus, we were able to identify which perturbations drove synchrony or asynchrony in this context and thus to explain their contribution using two simple metrics.

### For whom does dispersal matter? Importance of the biomass distribution

Knowing who disperses is crucial to understand biomass dynamics in metacommunities (Koelle and Vandermeer, 2005; Pedersen et al., 2016). However, even when dispersal is homogeneous among the various species (*i.e.* same dispersal rates *d*_*i*_ for all species), increasing dispersal does not affect all species in the same way (Fig.6). In fact, abundant species are more affected by demographic processes such as self-regulation, which scale as the square of biomass, or trophic interactions, which scale as the product of predator and prey biomass (see equation (18)). Thus, changes in dispersal rates lead to top-down or bottom-up coupling between patches depending on biomass distribution. Oncer we know for whom dispersal matters, the model can be simplified to a metacommunity where only a few species connect patches. With such a restricted dispersal, perturbing a species in one patch can lead to an opposite response in the other connected patch. In fact, perturbations affecting basal species have a bottom-up propagation (Fig.1C) and correlate all the species from the same food chain, while perturbations affecting top species have a top-down propagation and create trophic cascade correlation patterns (Fig.1D). Thus, if the perturbed species are not the dispersing species, both patches can display different correlation patterns, which can lead to anti-correlated responses of the different populations of the same species and hence to asynchrony between the different populations (Fig.A-1E, Fig.A-2A and A-2B in supporting information). The correlation or anti-correlation of populations depends on the shortest trophic distance from the dispersing species, as suggested by Wollrab et al. (2012). Species at odd distance have correlated population fluctuations, while species at even distance have anti-correlated population fluctuations (Fig.2A).

**Figure 6:**
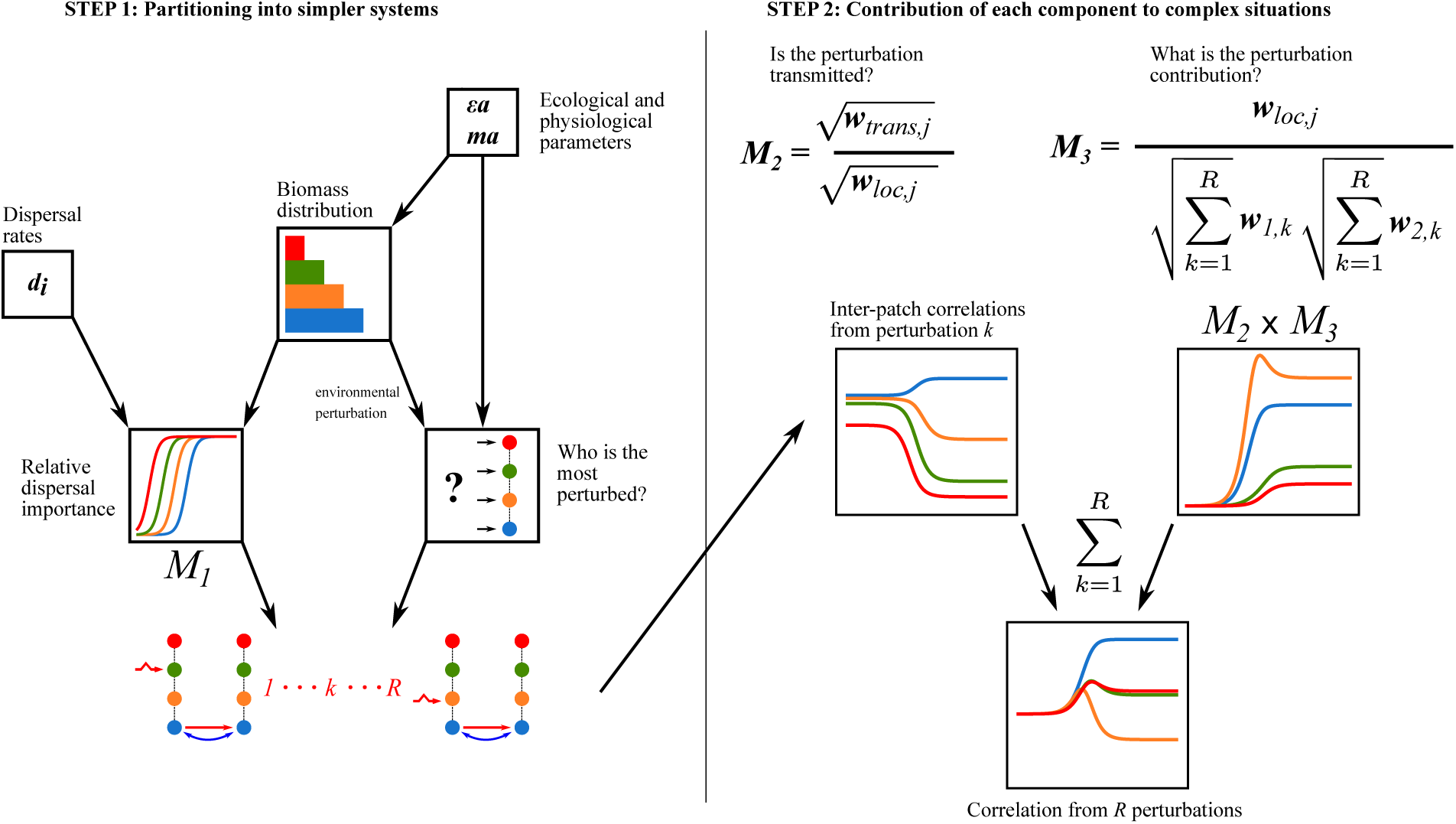
Sequential framework to understand the transmission of perturbations in meta-communities. Biomass distribution (driven by physiological and ecological parameters) is central as less abundant species are more affected by dispersal (metric *M*_1_) and are less affected by environmental perturbations. Knowing that, we can simplify the system into a metacommunity where only a few species disperse and are perturbed. The effects of each perturbations can then be partitioned to understand how much they contribute to the total correlation between patches. The contribution of each perturbation can be interpreted by two metrics: *M*_2_ that quantifies how much of the generated variability is transmitted through dispersal and *M*_3_ that quantifies the how much variability is generated compared to other perturbations. Therefore, *M*_2_ × *M*_3_ weights the correlation generated by each perturbation to reconstruct the correlation pattern obtained when multiple perturbations are applied.

The case where bottom-up perturbations are transmitted by top predators is related to the spillover process: a predator population thrives due to resource abundance in one patch and spills over to the other patches (Holt, 1984). For instance, favourable environmental conditions in the Baltic main basin increase cod abundance (bottom-up control) that colonise the Gulf of Riga, leading to a trophic cascade in this locality (top-down response) (Casini et al., 2012). More generally, predators cast a “shadow” that leads to trophic cascades around their source patch (McCoy et al., 2009). For instance, dragonfly that prey on flying insects around ponds reduce pollination there (Knight et al., 2005). Such dynamics of predators between natural habitats and crop fields are central in pest biocontrol (Tscharntke et al., 2012).

The bottom-up coupling between patches does not seem to be mediated by primary producers, which often have a low mobility (sessile terrestrial plants or drifting phytoplankton), but rather by non-living materials (Polis et al., 1997; Leroux and Loreau, 2008). Marleau et al. (2010) and Gounand et al. (2014) found in their models with limit cycles that flows of nutrient lead to anti-correlations between species populations, while we found a succession of correlations and anti-correlations. This suggests that systems with limit cycles respond differently to bottom-up coupling than systems in the vicinity of an equilibrium that receive stochastic perturbations because of processes such as phase-locking (Jansen, 1999; Liebhold et al., 2004; Vasseur and Fox, 2009). Abiotic resources can link very different food webs. For instance, mineral nutrients and dead organic matter link green and brown food webs (Wolkovich et al., 2014; Buchkowski et al., 2019) but additional mechanisms such as different food chain length, omnivory or stoichiometric constraints (Attayde and Ripa, 2008; Zou et al., 2016) make a direct comparison difficult. Nevertheless, our model gives basic insights into how a simple bottom-up coupling affects connected food chains dynamics and should improve our understanding of the additional effects brought by mechanisms such as different food chain length or stoichiometric constraints.

While top predator dispersal or basal resource diffusion have been extensively studied, the consequences of intermediate trophic level dispersal remain poorly understood. Our results show that the dispersal of intermediate trophic levels can dramatically change the correlation between populations of non-dispersing species. Pedersen et al. (2016) found that herbivores with a lower dispersal rate than primary producers or carnivores stabilise metacommunity dynamics (by having equilibria or asynchronous limit cycles). Most of the studies on coupled food webs considered systems displaying limit-cycles and largely ignored stochastic perturbations (McCann et al., 1998, 2005; Post et al., 2000; Koelle and Vandermeer, 2005). Our results suggest that dispersal patterns that leads to more asynchrony depends on which species is perturbed. If the most perturbed species is also the most affected by dispersal, it transmits the perturbation to all patches and synchronise them, thus reducing the stability of the system. Otherwise, asynchrony between patches can be promoted. Thus, the stabilising or destabilising effect of dispersal patterns is not absolute and depends on perturbations. In addition, perturbations can target specific species (*e.g.* harvesting, disease…) or affect all the species in different ways. For instance, Arnoldi et al. (2019) showed that environmental perturbations (*z* = 1) mostly affect abundant species (Fig.6 and see Fig.A-1B, A-2D and A-5 in the supporting information). Therefore, considering the biomass distribution is critical to fully understand the responses of coupled food chains to dispersal and perturbations.

### Multiple perturbation partitioning

Complex correlation patterns produced by multiple perturbations on different species in different patches can be easily partitioned into a sum of correlation patterns produced by a single perturbation (Fig.6). Such a partitioning is permitted by two characteristics of our model. First, the system is linearised. Thus, the temporal variations of each species in the vicinity of the equilibrium are the sum of the variations due to each interacting species. Second, the partitioning of the correlation pattern is permitted by the independence of the various perturbations. In fact, we can decompose the covariance matrix of perturbation *V*_*E*_ into a sum of matrices *V*_*Ej*_ corresponding to the perturbation of a single specie in a single patch (see equation (20) in supporting information). If some perturbations are correlated, we can still decompose the matrix *V*_*E*_ into a sum of independent blocks of correlated perturbations. The contribution of each perturbation in an assemblage of many independent perturbations can thus be easily understood as the product of the correlations between populations from the two patches weighted by the variability generated in each patch (Fig.6). Such a detailed partition of the contribution of each element of the system is not possible in systems displaying non-linear dynamics. For instance, Koelle and Vandermeer (2005) tested the effects of primary producers and top predator dispersal on population synchrony. They found that these two types of dispersal led to either asynchrony or synchrony between the populations of the other trophic levels but they were unable to go deeper in their interpretation. Their results are similar to our case where a perturbation is applied to top predators only and primary producers disperse (Fig.A-2B). Thus, the top predator-prey interaction must generate most of the variability in their system with limit cycles and may be equivalent to a perturbation of top predators in our linear system. Therefore, our model with linear dynamics could give clues to understand the response of models with non-linear dynamics.

Future investigations considering stochastic perturbations in models with type II functional responses are required to test the robustness of our results to non-linear dynamics. However, substantial modifications of our model are needed to make it consistent with previous models of couple food chains. In fact, simply considering type II functional responses does not lead to muti-trophic functional responses as in Post et al. (2000) or Rooney et al. (2006) when consumers disperse a lot and perfectly couple the two food chains. Independence between perturbations is also a key feature of our study as we explained earlier. Correlation between perturbations is expected to change the observed dynamics (Ripa and Ives, 2003; Vasseur and Fox, 2007). Leroux and Loreau (2012) considered reciprocal pulsed subsidies within a metacommunity model and demonstrated that the time delay between perturbations in each patch could reinforce or dampen the resulting oscillations. This suggests that the correlation pattern observed in our model when species from both patches are perturbed should be modified if perturbations are more or less correlated.

## Conclusion and perspectives

Our model demonstrates that species respond differently to dispersal depending on the biomass distribution in food chains even if all species disperse at the same rate. Thus, our simple model can serve as a good null model to test mechanisms involved in dispersal. Dispersal can be seen as a mechanism of optimal foraging where predator follow their prey in the patch where they are the most abundant. Dispersal also enables prey to escape their predators by migrating in a “refuge” patch where they are less abundant. This can be represented by density-dependent dispersal rates, which have a strong impact on dynamics (Hauzy et al., 2010; Liu et al., 2016). However, density-dependent dispersal changes the relative importance of dispersal and local demography as dispersal then scales with biomass similarly to self-regulation or predation, thus changing the interplay between dispersal and biomass distribution. Therefore, future studies should consider biomass distribution among species to properly assess the effects of dispersal on food chain dynamics.

When multiple perturbations are applied, the effects of each perturbation and each species can be partitioned in our model. Thus, future studies considering heterogeneity between patches would be able to isolate the contribution of the difference of parameters to food chain dynamics. For instance, Rooney et al. (2006) considered two food chains with different attack rates and coupled by a mobile top predator. In this case, perturbation partitioning would enable us to deeply understand how such differences between food chains may dampen perturbation transmission or promote asynchrony. Thus, our approach appears to be a promising tool to better understand the effects of many mechanisms that promote stability or asynchrony in coupled food chains or trophic metacommunities.

## Data accessibility

The C++ code of the simulations and the R code of the figures are available on Zenodo (doi:10.5281/zenodo.3613500).

## Appendix A Complementary results

### General responses of the food chain model to perturbations

**Figure A-1:**
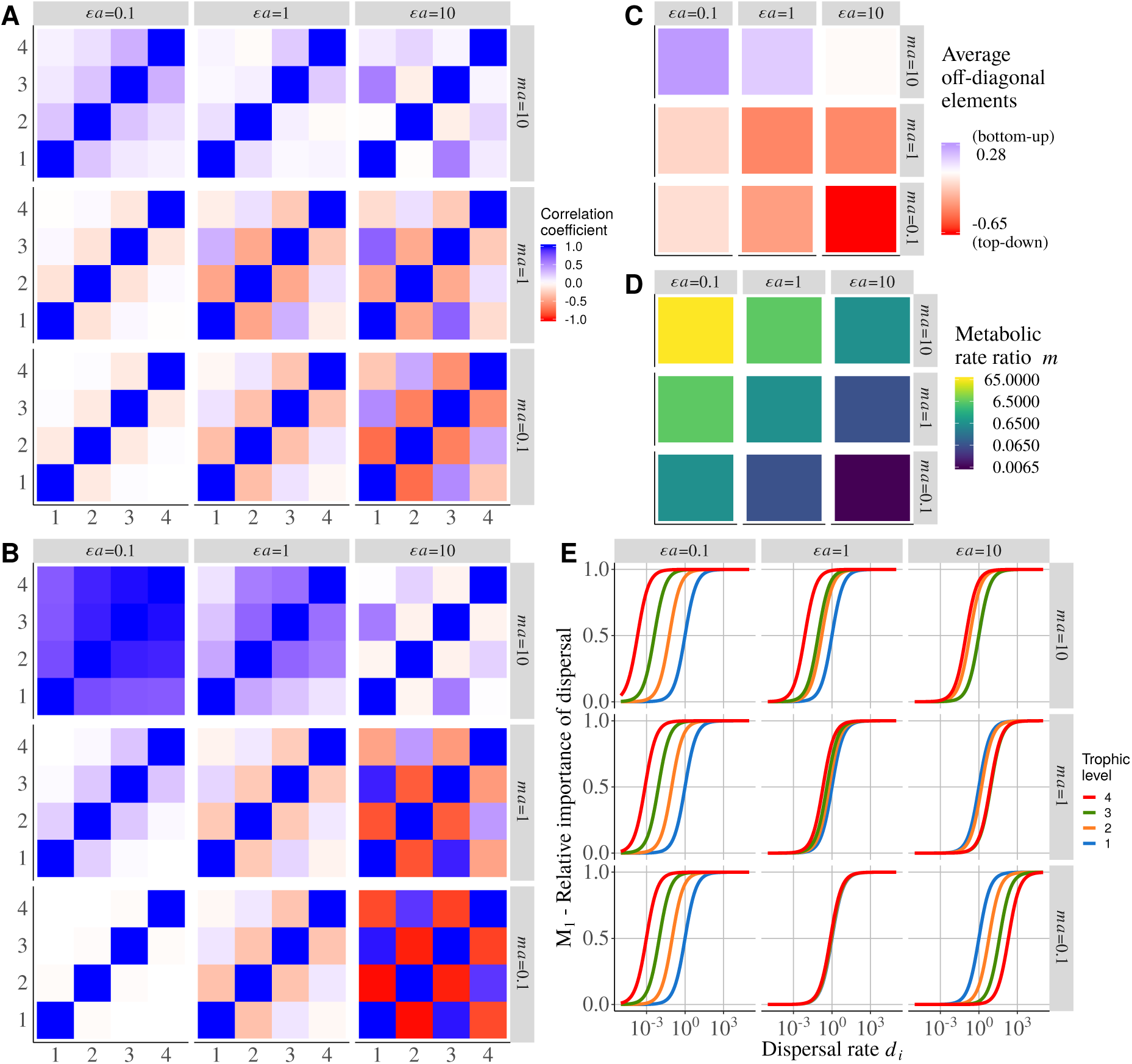
**A)** Correlation matrix within an isolated food chain (*d*_*i*_ = 0, no dispersal) and with demographic independent stochastic perturbations applied to each trophic level. Nine food chains with different combinations of *ϵa* and *ma* are tested. **B)** Same correlation matrix with environmental independent stochastic perturbations applied to each trophic level. **C)** General behaviour of food chains presented in **A**. All the off-diagonal elements are summed. If the output is positive, correlations are stronger and the food chain has a bottom-up response to perturbations, otherwise, anti-correlations are stronger and the food chain has a top-down response. **D)** Ratio of predator to prey metabolic rates (*m* = *m*_*i*_ + 1*/m*_*i*_). **E)** *M*_1_, ratio of dispersal process to the sum demographic and dispersal process (see equation (18)) for each trophic level along a dispersal rate gradient.

Fig.A-1 is complementary to Fig.1. In Fig.A-1A where all species receive independent demographic perturbations, adjacent trophic levels are correlated for *ma* = 10 while they are anti-correlated for *ma* ≤ 1 (bottom part of the graph). These two responses correspond respectively to bottom-up and top-down responses of the food chain to perturbations (Fig.A-1C). Such a difference is not due to biomass distribution among trophic levels as the variance of demographic perturbations is linear to species biomass. This is due to the values of *m* (= *m*_*i*+1_*/m*_*i*_, ratio of predator to prey metabolic rates, Fig.A-1D) that is much larger than 1 when the response of the food chain is strongly bottom-up and much lower than 1 when the response is strongly top-down. If predators have a faster metabolic rate than their prey, their demographic dynamics are fast and they recover from perturbations faster than their prey. This makes lower trophic levels relatively more sensitive to perturbations and leads to a bottom-up response to perturbations. The opposite situation makes predators more sensitive to perturbations and leads to a top-down response.

Environmental perturbations (*z* = 1, see equation (14), Fig.A-1B) lead to a distribution of bottom-up and top-down responses similar to Fig.A-1A but with stronger responses in the top-left and bottom-right corners of the graph. In fact, environmental perturbations affect more abundant species and are thus equivalent to demographic perturbations applied to a unique trophic level. Then, Fig.A-1B is a mix between Fig.1C and 1D depending on biomass distribution (Fig.1A).

Finally, *M*_1_, which is the relative importance of dispersal processes versus demographic processes (Fig.A-1E), completely depends on biomass distribution (Fig.1A): lower is the biomass, lower is the required dispersal rate *d*_*i*_ to affect species biomass dynamics.

### Transmission of perturbations to an undisturbed patch

The top-left corner (*ϵa* = 0.1 and *ma* = 10) of Fig.A-2A, where primary producers in patch #1 are perturbed, was detailed in Fig.3 but the same framework can be used to explain Fig.A-2A and Fig.A-2B, where top predators in patch #2 are perturbed, for all other combinations of *ϵa* and *ma*. If the perturbed species is also the most affected by dispersal (Fig.A-1E), then both patches are correlated as the perturbation is transmitted at the same trophic level in patch #2 as it is applied in patch #1 (for *ϵa* = 10 in Fig.A-2A and *ϵa* = 0.1 in Fig.A-2B). In other words, both patches receive almost the same perturbation at the same trophic level. In addition, the same pattern is observed in food chains where all species have the same biomass as they are equally affect by dispersal.

If the species sensitive to dispersal is not the perturbed one in patch #1, we observe a bottom-up response in one patch and a top-down response in the other one as each patch receives a perturbation at a different trophic level. This leads to the pattern detailed in Fig.5 and Fig.2. In Fig.A-2C where all species in patch #1 are perturbed, the top-left corner looks like Fig.A-2A and the bottom-right corner to Fig.A-2B. As isolated food chains have respectively bottom-up and a top-down responses in these two parts of the parameter space (Fig.A-1C), we can indeed expect responses similar to those where primary producers (Fig.A-2A) and top predators (Fig.A-2A) are respectively perturbed. Finally, Fig.A-2D, where all species in patch #1 receive environmental perturbations, is a mix between Fig.A-2A and Fig.A-2B as biomass distribution biases the effect of environmental perturbations towards the more abundant species. Thus, environmental perturbations in bottom-heavy biomass pyramids are equivalent to a perturbation of primary producers (top-left corner and Fig.A-2A) while in top-heavy biomass pyramids they are equivalent to a perturbation of top predators (bottom-right corner and Fig.A-2B).

**Figure A-2:**
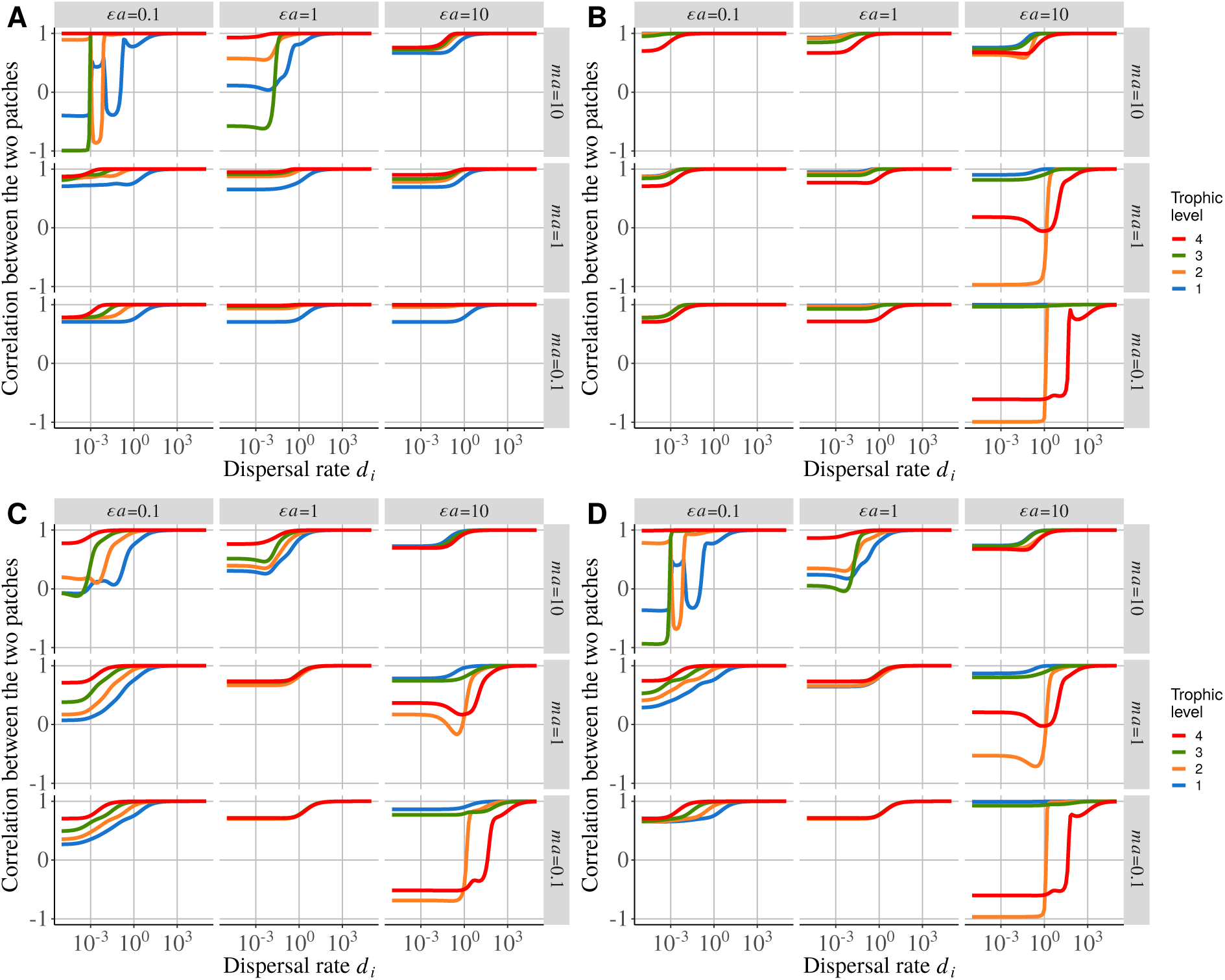
Correlation between populations of the same species from two patches with increasing dispersal rates (*d*_*i*_). All species are able to disperse. **A)** Only the primary producer of patch #1 receive a stochastic demographic perturbation. **B)** Only the top predator of patch #1 receive a stochastic demographic perturbation. **C)** All species of patch #1 receive independent stochastic demographic perturbations. **D)** All species of patch #1 receive independent stochastic environmental perturbations.

### Multiple perturbation partitioning

#### Mathematical demonstration of correlation partitioning

We consider *R* independent stochastic perturbations that can affect species from patch #1 or patch #2. Therefore, the covariance matrix of stochastic perturbations *V*_*E*_ is a diagonal matrix whose elements are equal to the variance 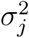 of each perturbation *j*. Thus, we define *V*_*Ej*_ the covariance matrix when only the *j*^*th*^ perturbation is applied and we have 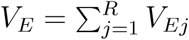.

From equation (16) we have:

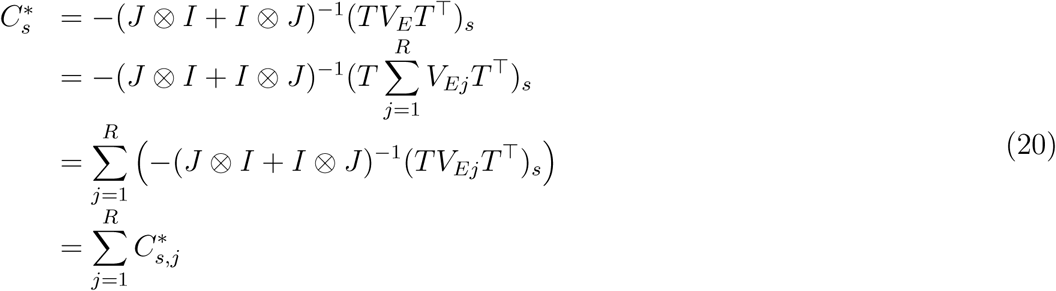

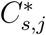 is the covariance matrix associated to *V*_*Ej*_. Thus, we have:

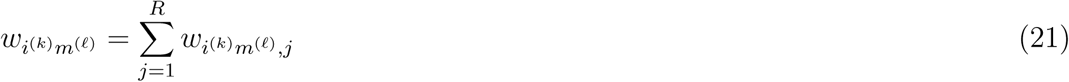

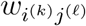 is the covariance between species *i* in patch *k* and species *j* in patch *ℓ* and is an element of the covariance matrix *C*\* and 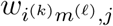 is an element of the covariance matrix 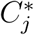. As we only consider two patches and are interested in the correlation between populations of the same species, we use the following notation:

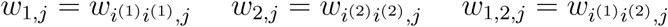

We also define *ρ*_*i*_ that is the correlation coefficient between the two populations of species *i* and *ρ*_*i,j*_ in the same correlation coefficient in the case where only the *j*^*th*^ perturbation is applied.

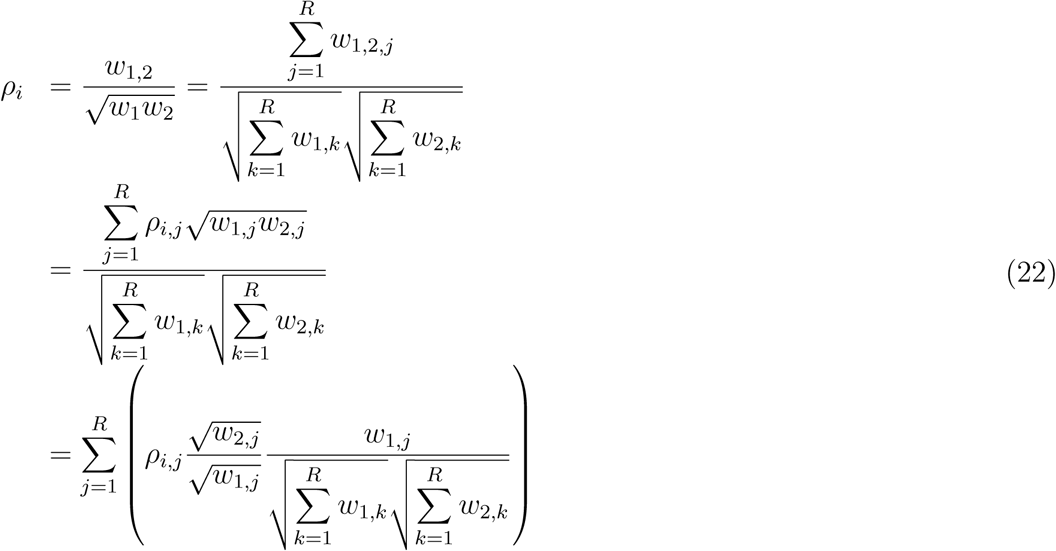

Eventually, the correlation pattern (*ρ*_*i*_) in a metacommunity where all species receive independent stochastic perturbations can be decomposed into a sum of the correlation patterns (*ρ*_*i,j*_) obtained when only one species is perturbed. The *ρ*_*i,j*_ are weighted by the variance generated in both patches by each perturbation.

#### Verification for the whole parameter space

The results found with a food chain sustaining four trophic levels (Fig.A-3) are consistent with the results from a food chain two with trophic levels (Fig.5). Primary producer populations become completely correlated as they are the only species able to disperse. Perturbing herbivores (Fig.A-3F), carnivores (Fig.A-3G) and top predators (Fig.A-3G) generates at least three times more variability in herbivore biomass than perturbing primary producers (Fig.A-3E). Thus, the average of the correlation patterns between herbivore populations in Fig.A-3B-D gives the moderated anti-correlation seen in Fig.A-3I and A-3J.

Carnivores are mostly and equally affected by the perturbation of carnivores (Fig.A-3G) and top predators (Fig.A-3H). Thus, averaging the corresponding correlation patterns (Fig.A-3C andA-3D) leads to the moderated correlation between the two carnivore populations (Fig.A-3I and A-3J).

Finally, top predators variability is mostly driven by their direct perturbation (Fig.A-3H), making the corresponding correlation pattern (Fig.A-3D) very similar to the correlation pattern obtained with multiple perturbations (Fig.A-3I and A-3J).

The partitioning of the effects of perturbations is also valid for the rest of the parameter space (Fig.A-4A and A-4B) and when only top predators are able to disperse (Fig.A-4C and A-4D) or when all species are able to disperse (Fig.A-4E and A-4F).

**Figure A-3:**
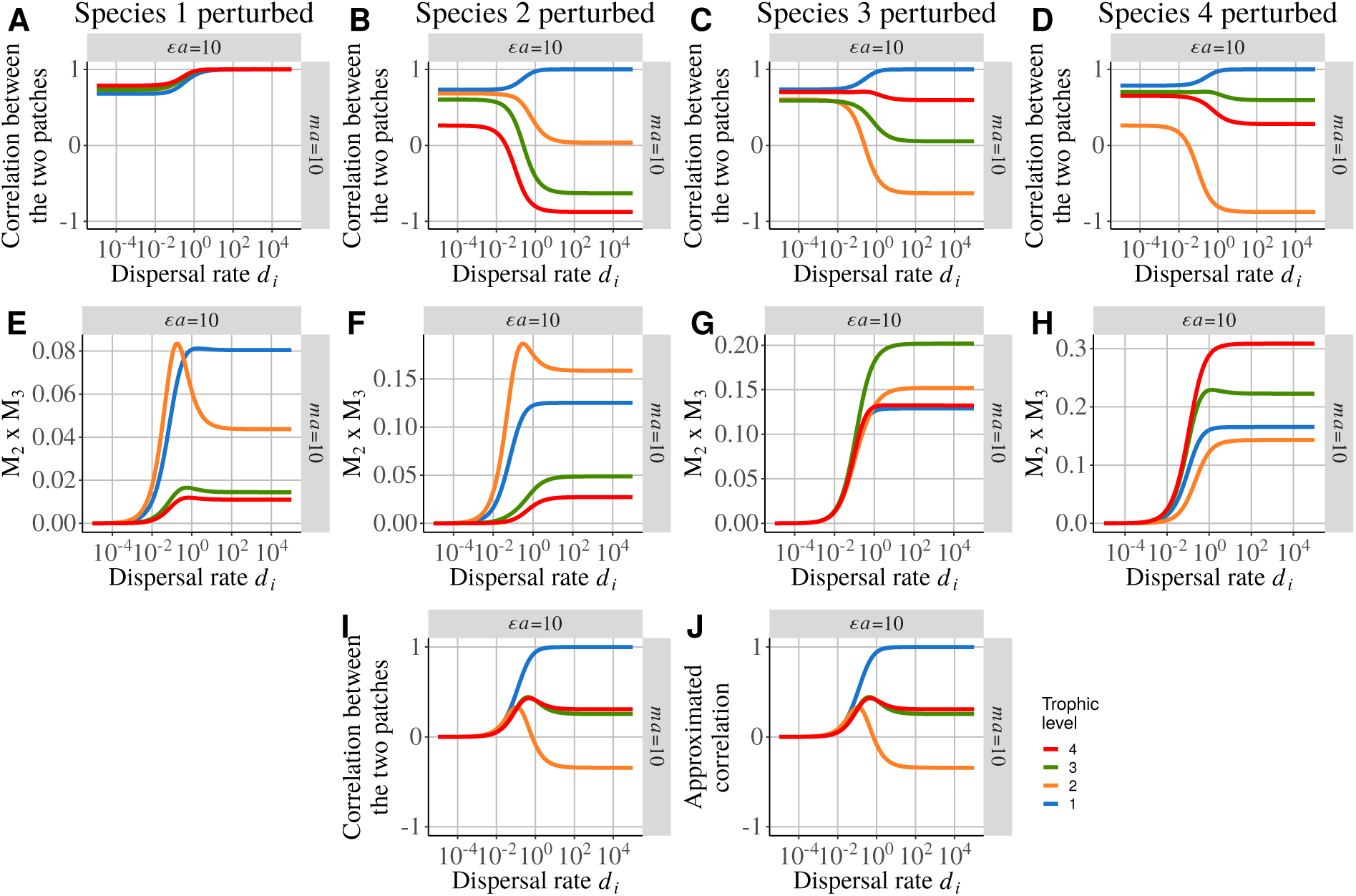
Detailed correlation pattern between two coupled food chains for *ϵa* = 10 and *ma* = 10 with increasing dispersal rates *d*_*i*_. Only primary producers are able to disperse. **A), B), C)** and **D)** are the correlation patterns between the two patches and **E), F), G)** and **H)** are the products of the two metric *M*_2_ and *M*_3_ when species 1, 2, 3 and 4 are respectively the only species perturbed. **I)** Correlation between patches when independent demographic stochastic perturbations are applied to all species in each patch. **J)** Reconstructed correlation pattern when all species in each patch are perturbed. It can by obtained as follow: **J**= 2×(**A**×**E**+**B**×**F**+**C**×**G**+**D**×**H**).

**Figure A-4:**
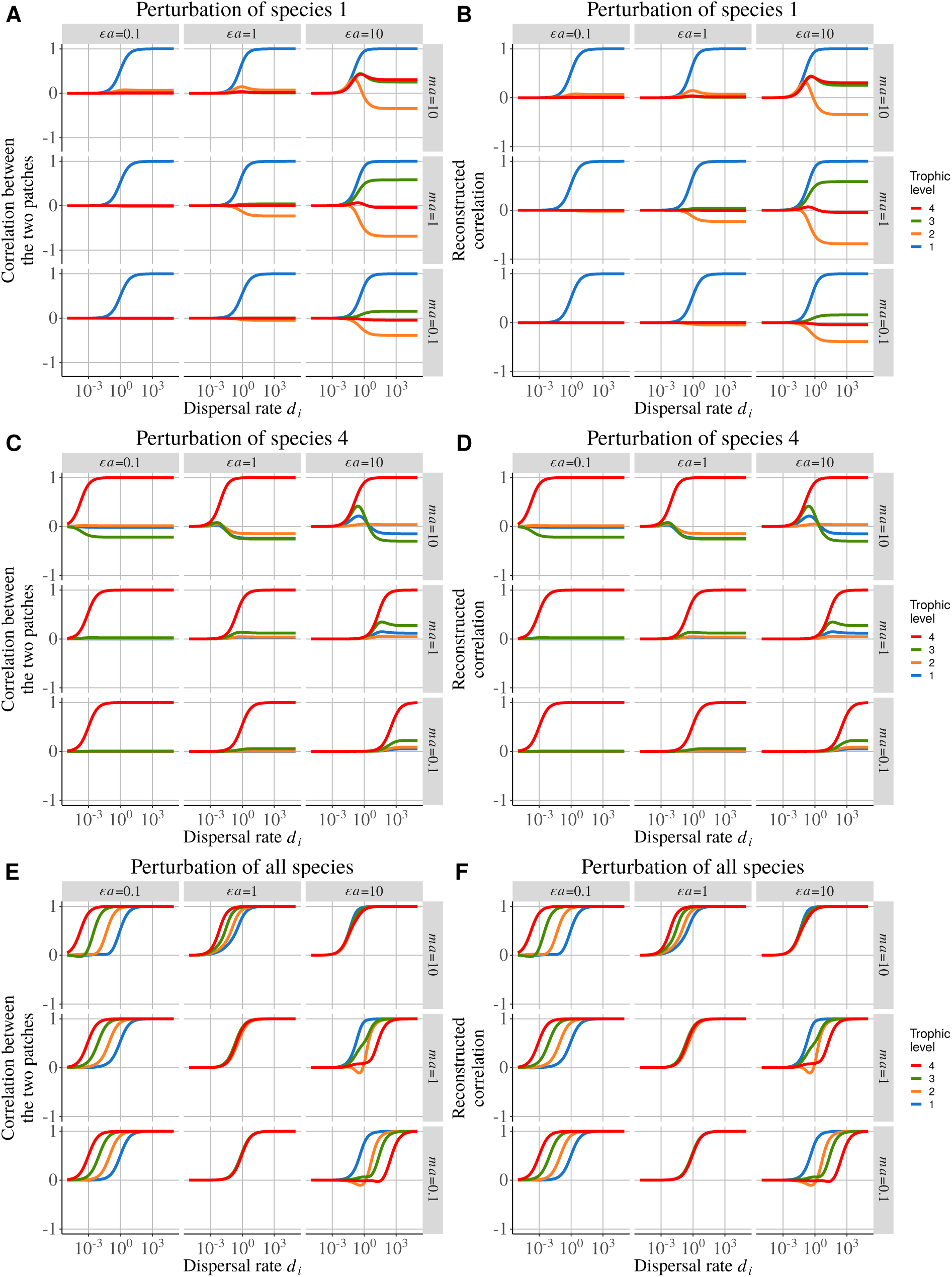
Correlation between populations of the same species from two patches with increasing dispersal rates *d*_*i*_. Independent demographic stochastic perturbations are applied to each species in each patch. **A)** Only primary producers are able to disperse. **C)** Only top predators are able to disperse. **E)** All species are able to disperse. Reconstructed correlation pattern when all species in each patch are perturbed. The reconstruction process is the same as explained in Fig.5 and Fig.A-3 but applied to the whole parameter space and for the cases where **B)** only primary producers **D)** only top predators are able to disperse and **F)** all species are able to disperse.

### Environmental perturbations

**Figure A-5:**
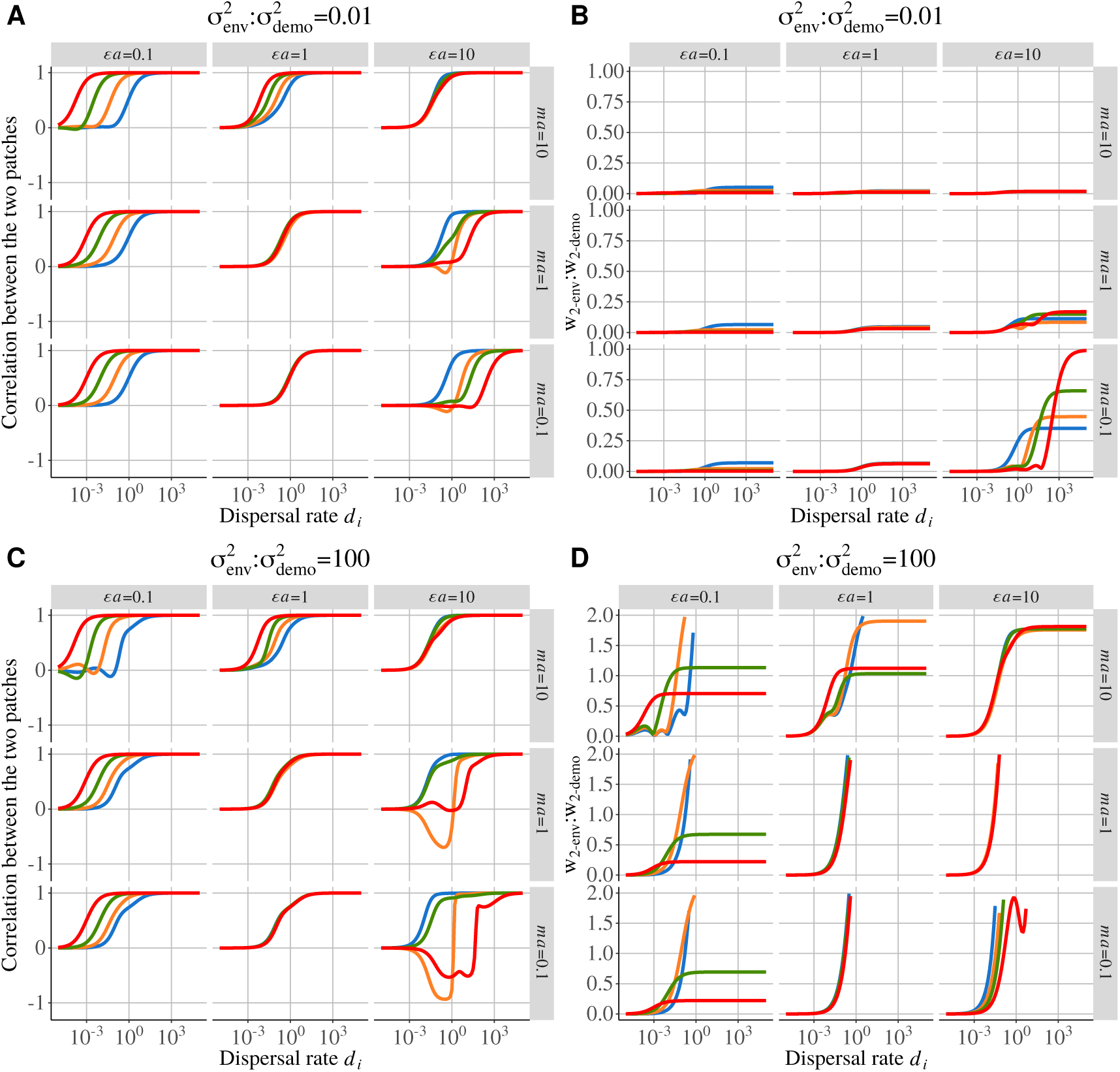
Correlation between populations of the same species from two patches with increasing dispersal rates *d*_*i*_. Independent demographic stochastic perturbations are applied to each species of each patch and independent environmental perturbations are applied to each species of patch #1. The relative weight of these two perturbations is given by the ratios of variance *w*_2−*env*_ in patch #2 when only species in patch #1 receive independent environmental perturbations to the variance *w*_2_−_*demo*_ in patch #2 when all species receive independent demographic perturbations. Two different ratio of environmental perturbation to demographic perturbation variance are tested: **A)** Correlation and **B)** biomass CV ratio for 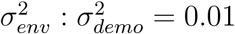 (stronger demographic perturbation) and **C)** Correlation and **D)** biomass CV ratio (y-axis cut for more readability) for 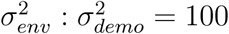 (stronger environmental perturbation).

Finally, we explore the effect of independent environmental perturbations (*z* = 1, see equation (14)) applied to all species in patch #1 in addition to demographic perturbations applied to all species in both patches. We test two ratios of environmental to demographic variances 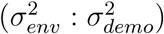 to understand how environmental perturbations can override the background demographic perturbations. Thus, we compare the variance generated in patch #2 when only environmental perturbations are applied in patch #1 to the variance generated when species from both patches receive independent demographic perturbations only.

Weak environmental perturbations 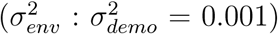 have no effects as Fig.A-5A is identical to Fig.A-4E (patch #1 and #2 receiving independent demographic perturbations only). At low dispersal rates, the variability due to the environmental perturbations transmitted to patch #2 through dispersal is completely overwhelmed by the variability due to demographic perturbations directly affecting species in patch #2 (Fig.A-5B). The increase in the transmitted variability from environmental perturbations at high dispersal rates is also jointed by the strong correlation of populations due to dispersal (Fig.A-2D). This increased correlation between patches due to dispersal overrides the increasing influence of environmental perturbations as all populations become completely coupled (bottom-right graph of Fig.A-5B).

However, strong environmental perturbations 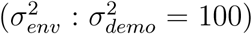 have dominant effects at intermediate dispersal rates as Fig.A-5C is similar to Fig.A-2D. At low dispersal rates, the variance ratio is low and patches remain uncorrelated (Fig.A-5D) while at high dispersal rate population are perfectly correlated. Food chains with top-heavy biomass pyramids (Fig.1A) seem particularly affected by environmental perturbations. Thus, changes of dispersal rate can dramatically switch population from anti-correlation to correlation.

## Appendix B Complementary material and methods

### Jacobian matrix

The Jacobian matrix *J* of the system can be expressed as:

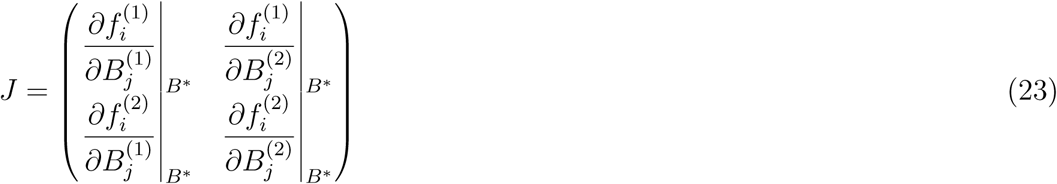

Where 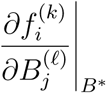 represents the effect of species from patch *ℓ* on dynamics of species from patch *k*. For simplicity, *J* can be split into blocks such as:

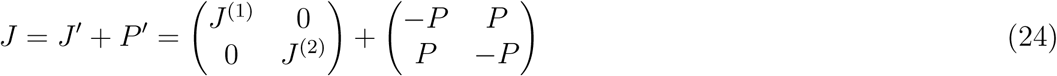

With *J*^(*k*)^ the Jacobian matrix of community *k* without dispersal, *P* the sub-dispersal matrix. *P* is defined by:

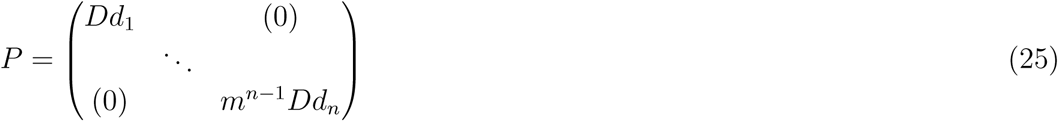

And *J* ^(*k*)^ by:

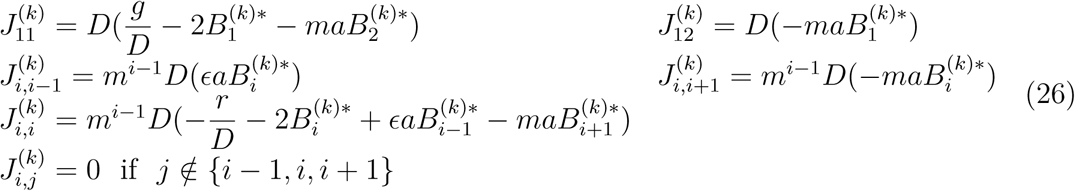

#### Parameters

**Table B-2:** Table of parameters. Only combinations of *m* and *a* leading to the desired values of *ma* were kept.

| | $\epsilon a = 0.1$ | $\epsilon a = 1$ | $\epsilon a = 10$ |
| --- | --- | --- | --- |
| $ma = 10$ | $m = 65$<br>$a = 0.15$ | $m = 6.5$<br>$a = 1.5$ | $m = 0.65$<br>$a = 15$ |
| $ma = 1$ | $m = 6.5$<br>$a = 0.15$ | $m = 0.65$<br>$a = 1.5$ | $m = 0.065$<br>$a = 15$ |
| $ma = 0.1$ | $m = 0.65$<br>$a = 0.15$ | $m = 0.065$<br>$a = 1.5$ | $m = 0.0065$<br>$a = 15$ |

